# Similarity in Spontaneous Tempo Impacts Flow Experience and Coordination Dynamics During Joint Performance

**DOI:** 10.1101/2025.09.16.676074

**Authors:** Leah Snapiri, Yoav Shapira, Noa Dagan, Ayelet N. Landau

## Abstract

Individuals tend to perform simple rhythmic tasks at a characteristic tempo, known as their spontaneous motor tempo. This motor trait, whether one naturally moves at a faster or slower pace, has been implicated in the ability to perform with external rhythmic structure across various tasks and experimental settings. In daily life, external rhythms often emerge through social interaction, as we continuously adapt to and move with rhythms generated by others. While previous studies have shown that individuals with similar spontaneous tempi can synchronize better in structured tasks, the role of individuals’ spontaneous tempo in unstructured, dynamic social interactions remains untested. In the current study, we examined how individuals’ spontaneous tempo shapes interpersonal coordination and subjective experience during an open-ended, free-form dyadic movement task. We first assessed individuals’ tempo using a spontaneous tapping task. Then, we measured interpersonal coordination patterns, including synchrony and leader-follower dynamics using a joint improvisation task inspired by the mirror game. Specifically, we developed a novel platform, composed of large transparent touch screens that resemble a shared window, on which individuals could continuously improvise motion together in an intuitive and playful manner. Participants alternated between game rounds with a designated leader and game rounds of leaderless improvisation. Following the mirror game, we also measured individuals’ subjective experience using a flow questionnaire. We found that individuals with similar spontaneous tempi reported greater flow experience during joint performance. Furthermore, we found that individuals with similar spontaneous tempi tended to synchronize **less** and were more variable and dynamic in their coordination patterns, specifically during leaderless improvisation rounds. Finally, we found that flow experience was highest when a participant’s movement tempo during the mirror game aligned with their own spontaneous tempo, regardless of their partner’s. These findings highlight individual spontaneous tempo as a meaningful trait that shapes both coordination dynamics and subjective experience during creative joint performance.

## Introduction

When performing together, individuals can enter states of optimal performance and experience, often referred to as moments of “togetherness”. These moments are characterized by effortless coordination, heightened arousal, and a shared sense of engagement and enjoyment (Noy et al., 2011; Noy, Levit-Binun, et al., 2015). What enables individuals to enter states of togetherness? Why are some encounters marked by seamless collaboration, while others remain effortful and disjointed?

In the present study, we explore how individual differences in spontaneous motor tempo shape interpersonal coordination and subjective experience during joint performance. Spontaneous motor tempo emerges naturally across a diverse range of motor tasks, such as tapping, walking, and speaking (Engler et al., 2024; Fraisse, 1982; Pfordresher et al., 2021). Previous research has shown that individuals vary in their spontaneous tempo (Desbernats et al., 2023; Fraisse, 1982; Hammerschmidt et al., 2021; McAuley et al., 2006), and that these tempo preferences are highly consistent over time, tasks, and measurement contexts (e.g. individual versus dyadic settings (Snapiri et al., 2025, unpublished manuscript), laboratory versus online environments, and musical versus non-musical tasks (Engler et al., 2024). Therefore, spontaneous tempo is a consistent motor trait that varies systematically across people.

Furthermore, individual differences in spontaneous tempo can also impact how people perform with external rhythmic structure in their environment. For example, McAuley and colleagues (2006) found that people are more accurate and stable when tracking auditory rhythmic stimuli near their spontaneous motor tempo. These findings extend to musical performance, where musicians perform better at tempi closely matching their spontaneous musical production tempo (Scheurich et al., 2018; Zamm et al., 2018). In the domain of temporal attention, we found that spontaneous motor tempo predicts perceptual facilitation within a rhythmic context in a visual discrimination task (Snapiri et al., 2023). Within a dyadic context, several studies have shown that individuals coordinate more effectively when they share similar tempo preferences (Alderisio et al., 2017; Snapiri et al., 2025, unpublished manuscript; Zamm et al., 2015, 2016). For example, Zamm and colleagues (2015, 2016) paired musicians based on their spontaneous production tempo and found that performance improved when partners had matching tempi. Similarly, Alderisio and colleagues (2017) found that groups with members that share similar individual movement frequencies synchronize better in joint rhythmic task compared to mixed groups. Extending this to non-rhythmic contexts, Słowiński and colleagues (2016) found enhanced coordination in dyads with similar velocities profiles during continuous joint movement. While this study focused on velocity rather than tempo, it highlights how aligning individual movement characteristics can facilitate interpersonal coordination. Together, these findings suggest that temporal coordination is optimized when individuals perform near their own tempo preferences or with partners who share similar movement characteristics.

While individuals with similar spontaneous tempo synchronize better during rhythmic, structured tasks, the role of tempo similarity in continuous, unstructured interactions remains largely untested. Previous research has focused on simplified coordination scenarios, often involving temporally or spatially structured tasks, where synchrony served as the primary measure of joint performance. However, more complex and naturally evolving interactions introduce various patterns of leading, following, and mutual adaptation that go beyond temporal matching. To capture these dynamics, we developed a novel experimental platform inspired by the mirror game paradigm. The mirror game paradigm is a dynamic improvisational task in which partners are instructed to move together, alternating between designated leader–follower roles and free improvisation (Noy et al., 2011). The mirror game offers a rich framework for examining interpersonal coordination as it unfolds over time. Furthermore, the mirror game has been shown to evoke moments of dyadic flow experience, often described as the dissolvement of self–other boundaries and the feeling of “being in the zone” (Noy et al., 2015). Therefore, it is a compelling paradigm to examine how spontaneous tempo shapes coordination dynamics and subjective experience while performing together.

To measure individuals’ spontaneous tempo, we used a simple tapping task, a widely used method for assessing spontaneous motor tempo (Engler et al., 2024; Fraisse, 1982; McAuley et al., 2006; Stern, 1900). We then invited pairs to play the MirrorPods™, a novel experimental platform developed in our lab consisting of two large transparent touch screens positioned face-to-face. Participants stood on opposite sides of the platform and moved together freely by sliding their index fingers across the transparent screens. This setup provided a natural and playful interactive space for joint movement, while enabling continuous recording of motion trajectories with high temporal and spatial resolution. We examined both overall synchrony and leader-follower dynamics across the duration of the interaction. Following the mirror game, participants completed the Activity Flow State Scale (AFSS, Payne et al., 2011), which captures subjective flow experience, including effortless attention and engagement (Nakamura & Csikszentmihalyi, 2014).

We hypothesized that similarity in spontaneous tapping tempo would impact movement synchrony and leader-follower dynamics between partners, as reflected by variations in temporal alignment over time. In addition, we hypothesized that greater similarity in spontaneous tempo would enhance individuals’ subjective experience of flow during joint performance. Finally, we examined how individual spontaneous tempo shapes the temporal structure of joint performance. Specifically, we characterized the overall frequency of individual motion segments within the continuous motion trajectory. We calculated the frequency separately for game rounds in which the individuals led the interaction, followed, or jointly improvised. We then asked whether individuals’ spontaneous tapping tempo is linked to the tempo preferences observed in their continuous, unstructured motion.

## Methods

### Participants

One hundred twenty individuals (90% right-handed, mean age = 23.66 [SD = 5]) participated in the experiment. All participants were female, as previous research has demonstrated stronger associations between nonverbal synchrony and subjective experience in female dyads compared to male dyads (Tschacher et al., 2014). Participants reported normal or corrected-to-normal vision, normal hearing, and intact motor abilities. Sixty-five percent of participants had no musical experience, while only seven and a half percent reported extensive musical training (10 years or more). Additionally, fifteen percent of participants indicated some experience with improvisation (1-4 years), with one participant reporting extensive improvisational experience (10 years). Dance experience was also documented, with six percent of participants reporting extensive dance training (10 years or more). All participants provided informed consent prior to the experimental session and received either monetary compensation or course credit for their participation. The experimental procedures received approval from the Ethics Committee of the Hebrew University of Jerusalem.

### Experimental Design

#### Assessing Spontaneous Tempo Preferences

Participants’ individual tempo preferences were assessed using the spontaneous tapping task. Participants were instructed to tap for one minute, at a regular tempo using the index finger of their dominant hand. The tapping task was performed twice, once on a small (10-inch) transparent touch screen and once on a large (55-inch) transparent touch screen. Both touch screens had a spatial resolution of <1 mm and a temporal resolution of 130 Hz. Tap times were extracted from the continuous touch signal using the Kivy library in Python (version 3.7). In the current study we focus on individuals’ spontaneous tempo as measured using the large transparent touch screen, which was used also for the dyadic part of the experiment. Overall, the two tapping measurements were highly correlated, *r* (117) =.87, *p* < .001.

#### Assessing Joint Performance

Joint performance was assessed using MirrorPods. MirrorPods are a custom build platform, composed of two 55-inch transparent touch screens. The screens are closely positioned to create the perception of a shared window through which participants can see each other and mirror each other’s movements (see Figure 1a). Participants were instructed to create motion together by moving their index fingers across the transparent touch screens. The position of each participant’s fingers was recorded at a temporal resolution of 125 Hz and a spatial resolution of <1 mm. The recorded x and y coordinates of each participant were extracted using Kivy library in Python (version 3.7) and synchronized in real time using the Lab Streaming Layer (LSL) software (Kothe et al., 2024).

**Figure 1.**
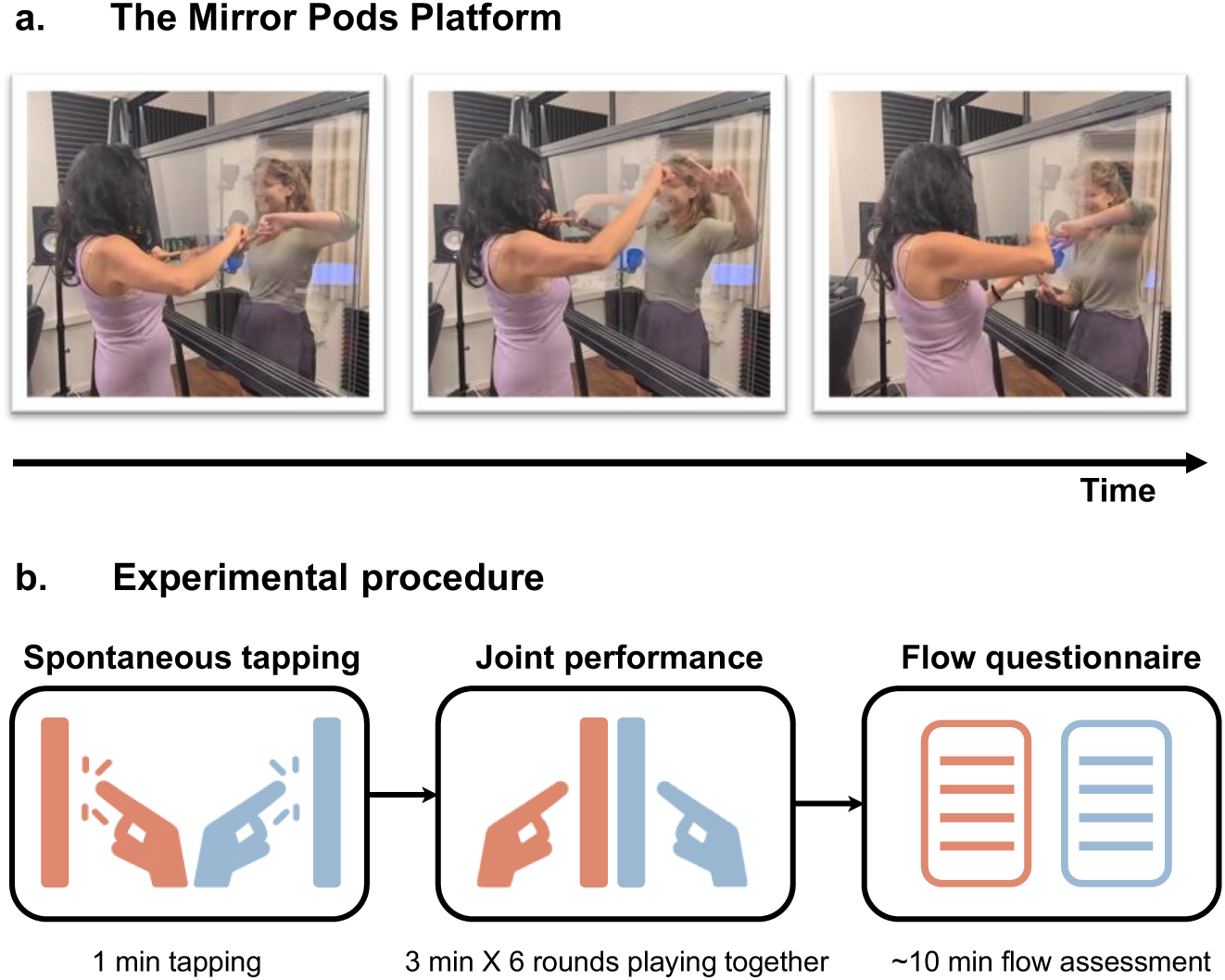
**(a) The Mirror Pods platform** consists of two closely positioned 55-inch transparent touchscreens. Participants stand facing each other on opposite sides of the setup and are instructed to move their index fingers across the screens. Finger positions are recorded at a temporal resolution of 125 Hz and a spatial resolution of less than 1 mm. Figure shows the authors (L.S. and N.D.) for illustrative purposes only; no participants are depicted. **(b) Experimental procedure.** Participants arrived at the lab in randomly assigned dyads. Each participant completed a one-minute spontaneous tapping task individually. Next, the dyads played the Mirror Game for six rounds of three minutes each, alternating between game rounds with a designated leader and game rounds of joint improvisation. Participants were instructed to move together in a synchronized manner that is fun and interesting for them. At the end of the session, each participant completed a self-report flow questionnaire assessing their subjective experience during the joint activity.

#### Assessing Subjective Experience

Subjective experience was assessed using the Activity Flow State Scale (AFSS, Payne et al., 2011). The AFSS is designed to measure the state of flow during physical activity, as in the context of physical rehabilitation or sports. It is composed of 26 items that are rated on a 5-point Likert scale. The items assess core flow dimensions, such as loss of self-consciousness, effortless focus and enjoyment. Example items include*: “I wasn’t concerned with how I was presenting myself”, “I had total concentration”,* and *“The experience left me feeling great”*.

### Procedure

Participants arrived at the lab in pre-assigned dyads and provided written informed consent. After completing a personal information questionnaire, they performed the spontaneous tapping tasks individually. After both participants completed the tapping task, they proceeded to the mirror game. Standing on either side of the MirrorPods, they used their index fingers to create synchronized and interesting motion together in three-minute rounds. In the leader-follower rounds, one participant led the interaction while the other followed. In the joint improvisation rounds, they created motion together without a designated leader. Each round began with a brief practice period to clarify instructions. Participants played a total of six rounds. The first three followed a fixed sequence: (1) Leader–Follower (L-F), (2) Follower– Leader (F-L), and (3) Joint Improvisation (JI). The final three rounds repeated these conditions in a counterbalanced order, ensuring that any observed differences could not be attributed to the sequence of play. After the mirror game, participants completed the AFSS questionnaire (see Figure 1b).

## Data Analysis

### Preprocessing and Outcome Measures

#### Spontaneous Tempo

To estimate each participant’s spontaneous tapping tempo, we calculated the mean inter-tap interval (ITI) between successive taps. Prior to computing the mean, we excluded extreme ITIs, that were more than 1.5 interquartile ranges above the upper quartile or below the lower quartile of the participant’s ITI distribution (on average 2.9% (SD = 3.38)). Then, for each dyad, we computed the **spontaneous tempo distance**, defined as the absolute difference between the participants’ mean ITIs. We also calculated the **dyadic tempo**, defined as the average spontaneous tapping tempo of the two participants. Six dyads were removed from farther analysis as their spontaneous tempo distance exceeded three median absolute deviations (MAD) from the median spontaneous tempo distance.

#### Joint Performance

We first filtered the position trace of each finger using a fourth-order, two-way low-pass Butterworth filter with a 5 Hz cutoff frequency (Noy et al., 2015), applied separately to the X and Y coordinates. To convert the relative touchscreen values to physical units, the X and Y coordinates were scaled according to the screen’s dimensions (width: 1250 mm, height: 720 mm). Based on the filtered X and Y coordinates, we then calculated the finger’s distance from the origin (the bottom-left corner of the screen) on a sample-by-sample basis. Specifically, the distance at each time point, *d(t)*, was computed as: 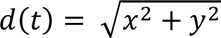, where *x* and *y* represent the filtered coordinates of each finger. This transformation allowed us to capture each finger’s movement trajectory as a one-dimensional time series.

To quantify the quality of interpersonal coordination we used two complementary measures: zero-lag synchrony and coordination variability. Both measures were computed using windowed cross-correlation, as implemented in the corrgram function in MATLAB (Marwan, 2025). We used windowed cross-correlation to address the non-stationary nature of our motion data (Boker et al., 2002; Konvalinka et al., 2010). Specifically, we computed the cross-correlation function between the two motion trajectories using a moving window of 4 seconds, a step size of 2 seconds, and a maximum lag of 2 seconds. For each window, we extracted **(a)** the correlation values at zero-lag, and **(b)** the time lag at which the two motion traces reached maximal cross-correlation.

##### Zero-lag synchrony

To obtain a single synchrony score for each coordination condition we averaged the correlation values at zero lag across all time-windows within each game round. Then, the synchrony values were averaged across the two fingers and across the two repetitions of each condition (LF, FL, JI). Higher synchrony scores reflected greater moment-to-moment alignment between participants’ movement trajectories.

##### Coordination variability

To quantify fluctuations in temporal alignment over time, we calculated the variability in the time lags at which maximal cross-correlation occurred across the game duration. For each time window, the lag corresponding to the peak cross-correlation value indicated both the direction of coordination, that is, which participant was leading (positive lags reflect one participant leading, negative lags the other), and the magnitude of the temporal offset between the participants. We calculated the standard deviations of the distribution of the time lags across all windows within each game round. Then we averaged the standard deviation values across the two fingers and the two repetitions of the same coordination condition. Higher values indicated more dynamic and variable alignment over time, whereas lower values reflected more stable and predictable coordination dynamics.

#### Spontaneous Tempo Preferences in Joint Performance

In addition to calculating coordination measures we also estimated the temporal structure of joint performance. To this end we segmented the continuous motion trajectories into elementary motion strokes, defined as a period of movement between adjacent zero-crossings in the velocity trace (Hart et al., 2014; Noy et al., 2011). Namely, between points in time when individuals changed the movement direction (moving either toward or away from the origin). We excluded segments that lasted below 50 milliseconds, or if the total distance traversed during the segment was less than 2 mm. Then, to extract individual tempo preferences during joint performance we calculated the number of segments per second (SPS), for each participant, as a leader, a follower, and during joint improvisation. Specifically, we divided the overall number of produced segments by the total duration of the recording to capture global tempo characteristics.

Due to technical issues, the motion data of 4 out of 60 dyads was not recorded. Additionally, during initial stages of data collection some dyads had insufficient touch recordings during specific game rounds. To ensure measurement reliability, we excluded game rounds containing less than 20% of the expected recording duration. This resulted in the exclusion of 8.9% (30/336) of recordings. In addition, 2 dyads were removed due to abnormal task performance (lack of movement/ Erratic movement).

**Figure 2.**
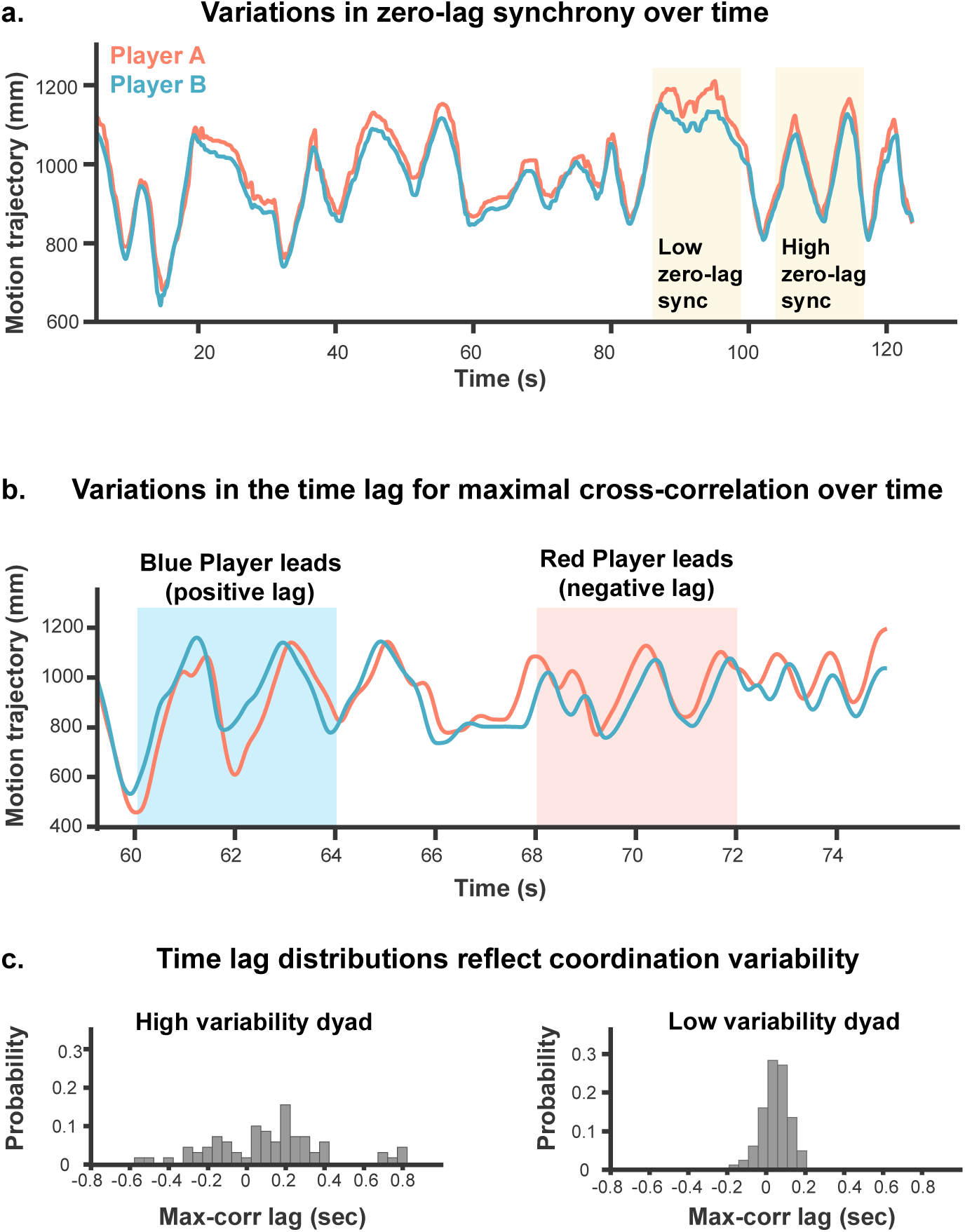
**(a) Example of fluctuations in zero-lag synchrony over time**. Movement trajectories of one finger from each participant (red: Player A, blue: Player B) are presented for a single dyad during a segment of joint improvisation. Movement trajectories are represented by the distance of the finger (in mm) from the origin point (bottom left corner of each screen). Yellow rectangles highlight periods of relatively high and low zero-lag synchrony. **(b) Example of a shift in leader–follower roles.** We present several seconds of joint improvisation of a single dyad. For each 4-second window we extract the time lag at which maximal cross-correlation occurs. In the first window (blue highlight), Player B precedes Player A. This results in a positive lag between the two players as Player B’s movement must be shifted forward in time to best align with Player A. In the second window (red highlight), Player A (red line) precedes. This results in a negative lag, as Player B’s movement must now be shifted backward to achieve optimal alignment. **(c) Two example dyads illustrating different coordination variability scores.** Coordination variability is quantified as the standard deviation of lag values extracted from overlapping 4-second windows throughout the entire game duration. The left panel shows a lag values distribution of a dyad with high coordination variability. The distribution is characterized by both positive and negative lags and a widespread, indicating a dynamic and less predictable coordination pattern. The right panel shows a dyad with low coordination variability, where lags are narrowly centerer close to zero, indicating stable and predictable temporal alignment over time.

#### Subjective Experience

To quantify participants’ flow experience during the mirror game, we calculated their total score on the AFSS as a percentage of the maximum possible score (130 points). If a participant missed an item (7/120 participants), their percentage was computed based on the total number of items they answered. No participant missed more than one item. To obtain dyadic flow, we averaged the individual flow scores of both participants within each dyad. We removed one dyad from further analysis due to language barriers that interfered with their ability to complete the questionnaire.

### Statistical Analysis

#### Variable coding

Coordination conditions were re-coded based on the relative spontaneous tempo of the leader in each round. Specifically, Leader–Follower and Follower–Leader rounds were categorized as either Fast Lead or Slow Lead, depending on whether the leader had a faster or slower spontaneous tempo than the follower. The resulting coordination condition variable had three levels: Fast Lead (FL), Slow Lead (SL), and Joint Improvisation (JI). This allowed us to examine whether performance is modulated based on the relative spontaneous tempo of the leader within each dyad.

#### Primary Analyses

To assess the impact of spontaneous tempo distance on synchrony and coordination variability across the three coordination conditions, we used linear mixed-effects models as implemented in the ‘lmerTest’ package in R version 4.2.2 (Kuznetsova et al., 2017). The models included spontaneous tempo distance, dyadic tempo, coordination condition (Fast Lead, Slow Lead, Joint Improvisation), and their interactions as fixed effects, and random intercepts for dyads to account for the nested structure of the data. Coordination condition was dummy coded with Joint Improvisation as the reference level. All continuous measures were centered and scaled. Post-hoc comparisons were performed using the ‘emmeans’ package in R (Lenth, 2019), and *p*-values were FDR-corrected where appropriate (Benjamini et al., 2001).

To assess how individuals’ spontaneous tempo relates to the temporal structure of joint movement, we conducted separate linear regressions predicting segments per second (SPS) from spontaneous tapping tempo in each coordination condition. In a follow-up analysis, we examined whether alignment between individuals’ spontaneous tapping tempo and their movement tempo during joint performance predicted flow experience, using the interaction terms in a multiple regression analysis.

#### Outlier removal

Prior to statistical analysis, we excluded dyads or individuals whose outcome measures exceeded three median absolute deviations (MADs) from the median within each experimental condition (Fast Lead, Slow Lead, and Joint Improvisation for dyadic-level analyses; Leading, Following, and Joint Improvisation for individual-level analyses). This procedure resulted in the removal of 4.24% of dyads on average per measure and condition at the dyadic level (SD = 2.26%) and 2.94% of participants on average at the individual level (SD = 1.70%). Additionally, one dyad was excluded from all flow-related analyses due to an extreme flow score that exceeded the outlier threshold.

## Results

### Spontaneous Tempo Similarity Enhances Flow Experience during Joint Performance

To examine the influence of spontaneous tempo on dyadic flow experience, we conducted a multiple linear regression analysis with spontaneous tempo distance, dyadic tempo, and their interaction as predictors. The dependent variable, dyadic flow, was calculated based on the average flow ratings reported by both members of each dyad. The overall model was statistically significant, *F* (3, 48) = 4.74, *p* = .006, accounting for 18% of the variance in flow scores (adjusted *R*² = .18). Specifically, enhanced flow experience was linked to decreased spontaneous tempo distance, *β* = −0.49, 95% CI [−0.76, −0.23], *t* (48) = −3.75, *p* < .001, indicating that dyads with similar spontaneous tempo reported greater levels of flow (see Figure 3).

**Figure 3.**
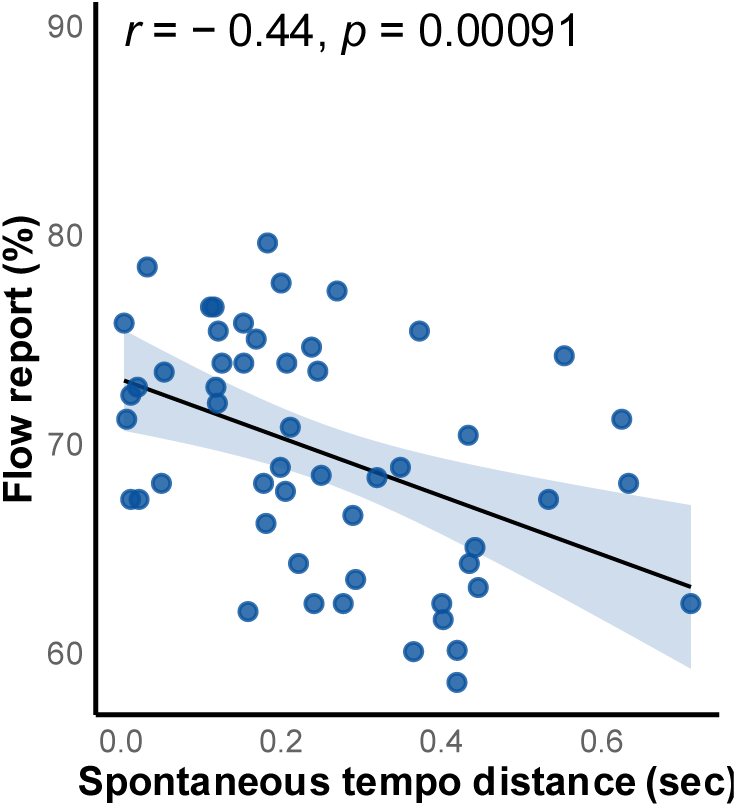
Dyads with more similar spontaneous tempi reported higher flow experience during joint performance. The x-axis represents the absolute difference in spontaneous tempo between partners (in seconds), and the y-axis shows the dyadic flow score (in percent). Greater tempo distance between partners was associated with lower reported flow.

Dyadic tempo, defined as the average spontaneous tempo across both members of the dyad, did not impact flow experience, *β* = 0.01, 95% CI [−0.26, 0.28], *t*(48) = 0.08, *p* = .938, nor did the interaction between dyadic tempo and spontaneous tempo distance, *β* = −0.20, 95% CI [−0.62, 0.23], *t*(48) = −0.93, *p* = .356. Therefore, the absolute tempo of a dyad, whether relatively fast or slow, does not impact participants’ flow experience.

### Spontaneous Tempo Similarity is Associated with Less Synchrony During Joint Performance

To examine the impact of spontaneous tempo on synchrony during joint performance, we conducted a mixed-effects multiple regression analysis with spontaneous tempo distance, dyadic tempo, coordination condition (Fast Lead, Slow Lead, Joint Improvisation), and their interactions as fixed effects. We also included random intercepts for each dyad to account for the nested data structure. Synchrony was operationalized as the mean zero-lag correlation between participants’ motion trajectories. The model explained a substantial proportion of the variance in synchrony scores, with a conditional *R*² of .54. The fixed effects alone accounted for 20% of the variance (marginal *R*² = .20).

As shown in Figure 4a, having a designated leader did not significantly affect overall synchrony, both when comparing joint improvisation to rounds where the relatively fast member of the dyad led the interaction, *β* = −0.19, 95% CI [−0.52, 0.14], *t*(116) = −1.15, *p* = .254, and when comparing joint improvisation to rounds in which the relatively slow member led the interaction, *β* = −0.16, 95% CI [−0.49, 0.17], *t*(116) = −0.95, *p* = .345. Furthermore, whether a dyad consisted of relatively fast or slow individuals in their spontaneous tempo did not impact synchronization, *β* = −0.04, 95% CI [−0.34, 0.25], *t*(116) = −0.28, *p* = .777.

**Figure 4.**
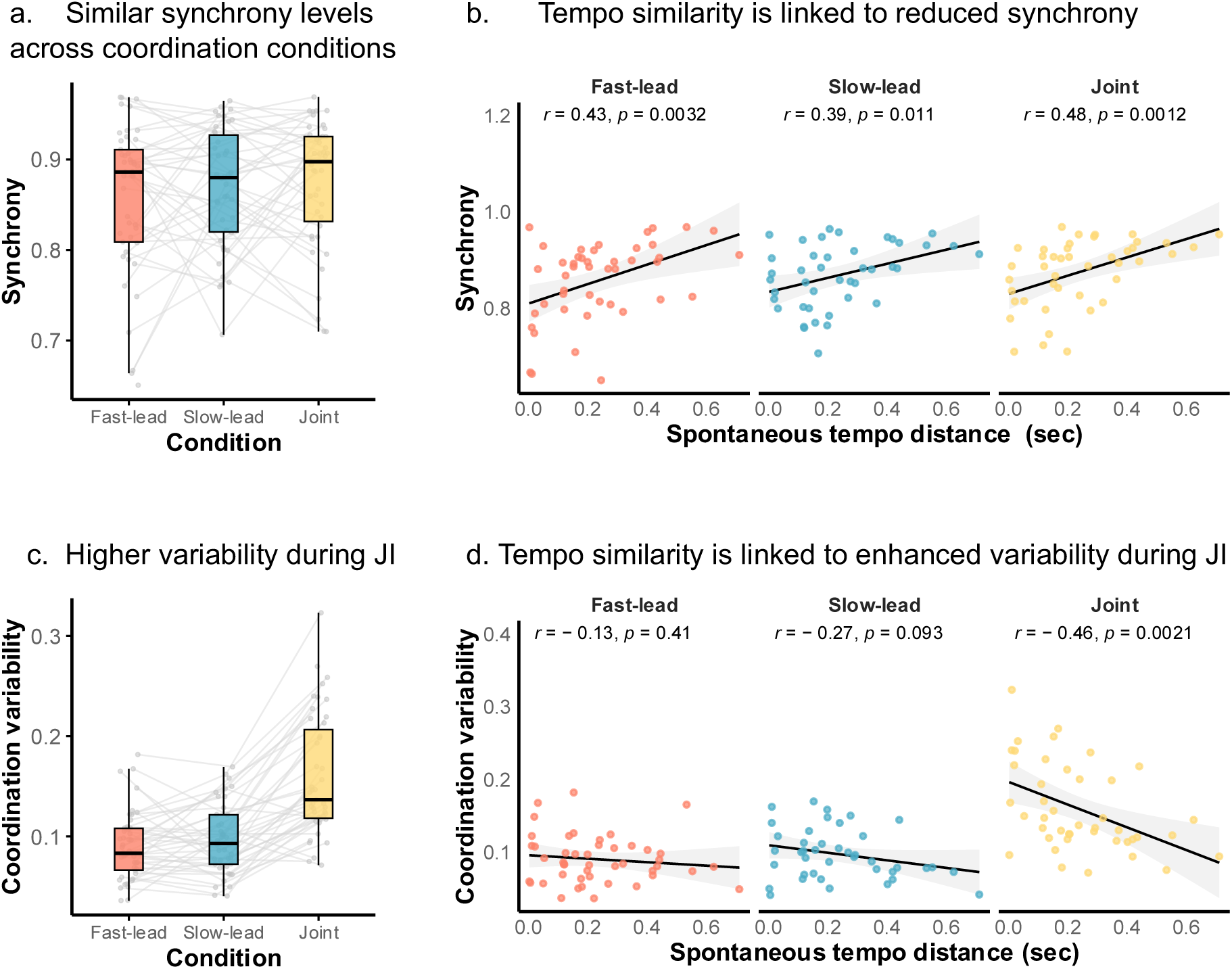
**(a) Mean zero-lag synchrony across coordination conditions**. Participants showed similar levels of synchrony across the fast-lead, slow-lead, and joint improvisation conditions. Dyadic synchrony was quantified as the average zero-lag cross-correlation across overlapping 4-second windows within each condition. **(b)** Tempo similarity is linked to reduced synchrony. We plot the relationship between spontaneous tempo distance (x-axis, in seconds) and zero-lag synchrony (y-axis), separately for each condition. Across all conditions, dyads with more similar spontaneous tempi (i.e., smaller tempo distance) exhibited **less** synchrony. **(c) Enhanced coordination variability during joint improvisation rounds.** Coordination variability was calculated based on the standard deviation of the time lags at which participants achieved maximal cross-correlation. These lag values were extracted from overlapping 4-second windows across the entire game. Dyads exhibited significantly greater coordination variability during joint improvisation compared to both fast-lead and slow-lead conditions, suggesting more dynamic and less predictable coordination patterns in the absence of a designated leader. **(d) Tempo similarity is associated with enhanced coordination variability, specifically during joint improvisation.** We plot the relationship between spontaneous tempo distance (x-axis, in seconds) and coordination variability (y-axis), separately for each condition. While tempo distance was unrelated to variability in the fast- and slow-lead conditions, during joint improvisation, dyads with similar spontaneous tempi (i.e., smaller tempo distance) showed **greater** coordination variability, indicating more dynamic and variable temporal alignment over time.

However, the distance between the spontaneous tempi of the individuals within each dyad was significantly correlated with dyadic synchronization, *β* = 0.42, 95% CI [0.13, 0.71], *t* (116) = 2.85, *p* = .005. Specifically, individuals with similar spontaneous tempi were less synchronized in their motion trajectories across all conditions. This was reflected in a positive relationship between spontaneous tempo distance (larger values reflect individuals with more dissimilar spontaneous tempi) and synchrony as shown in Figure 4b (**Fast Lead**: *r* (43) = .43, *p* = .003, **Slow Lead**: *r* (40) = .39, *p* = .010, **Joint Improvisation**: *r* (42) = .48, *p* = .001).

### Spontaneous Tempo Similarity is Associated with More Variable Coordination Dynamics

We next examined the impact of spontaneous tempo distance, dyadic tempo, and condition on coordination variability. Coordination variability was calculated based on the standard deviation of the time lags at which individuals reached maximal cross correlation. High coordination variability scores reflect dynamic and unpredictable coordination patterns. Low coordination variability scores reflect stable temporal alignment over time. The full model explained 66% of the variance in coordination variability (*conditional R*² = .66), with 44% attributable to the fixed effects (*marginal R*² = .44). As can be seen in Figure 4c, individuals exhibited enhanced variability in their coordination patterns during joint improvisation rounds compared to rounds with a designated leader. This tendency was observed both when comparing joint improvisation to Fast Lead rounds (*β* = −1.34, 95% CI [−1.62, −1.06], *t* (116) = −9.51, *p* < .001) and to slow Lead rounds (*β* = −1.16, 95% CI [−1.45, −0.88], *t* (116) = −8.07, *p* = .001). This pattern is expected, as joint improvisation involves more flexible turn-taking, resulting in both positive and negative time-lags between the two participants, whereas designated leader conditions exhibit more unidirectional time lags. In addition, we found that individuals with similar spontaneous tempi were more variable in their coordination pattern, *β* = −0.55, 95% [−0.79, −0.31], *t* (116) = −4.59, *p* < .001. This relationship between tempo similarity and coordination variability was significantly modulated by the experimental condition. Specifically, the association between tempo similarity and variability was strongest during joint improvisation compared to leader-follower conditions (**Fast Lead vs. Joint Improvisation**: *β* = 0.46, 95% CI [0.20, 0.72], *t*(116) = 3.45, *p* < .001, **Slow Lead vs. Joint improvisation**: *β* = 0.33, 95% CI [0.06, 0.6], *t*(116) = 2.45, *p* = .02). We also conducted a Pearson correlation analysis to directly examine the relationship between spontaneous tempo distance and coordination variability within each condition (Figure 5). These analyses confirmed that smaller tempo distance was linked to enhanced coordination variability specifically during joint improvisation (**Fast Lead**: *r* (44) = −.13, *p* = .41 **Slow Lead**: *r* (39) = −.27, *p* = .09; **Joint Improvisation**: *r* (41) = −.46, *p* = .002. figure 5). The impact of dyadic tempo on coordination variability did not reach significance, *β* = 0.19, 95% CI [−0.05, 0.44], *t* (116) = 1.55, *p* = .12, All other interaction were not significant.

**Figure 5.**
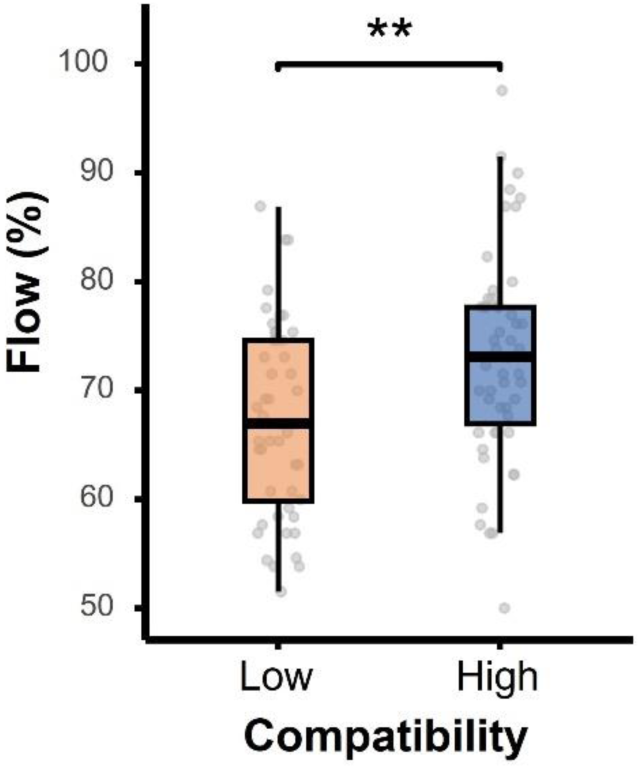
High tempo compatibility enhances flow experience. Participants were divided into two groups based on a tempo compatibility index, which reflects the alignment between standardized scores of spontaneous tapping tempo and the tempo of joint performance (SPS). A **low compatibility index** indicates that the two scores fell on opposite sides of the distribution (e.g., a fast tapper with a slow joint tempo, or vice versa), while a **high compatibility index** indicates that both scores were on the same side (e.g., both fast, or both slow). Asterisks indicate significance at **p < .01**.

### Tempo Preferences in Continuous Motion When Leading, Following and Jointly Improvising

To examine the link between spontaneous tapping tempo and the temporal structure of joint performance, we conducted a linear regression analysis with spontaneous tapping tempo as the predictor and SPS (segments per second) as the outcome measure. Analyses were performed separately for rounds in which the participant led the interaction, followed, or jointly improvised. After correcting for multiple comparisons, none of the tested links remained significant. Therefore, overall, individuals’ spontaneous tapping tempo did not predict the tempo of joint performance.

### High Compatibility Between Dyadic Performance Tempo and Individuals’ spontaneous Tempo Results in Enhanced Flow Experience

Given that the temporal structure of joint performance was not determined by individuals’ spontaneous tempo preferences we next examined how the compatibility between individuals’ spontaneous tapping tempo and joint performance tempo affects subjective experience. We conducted multiple linear regression analyses with spontaneous tapping tempo, SPS, and their interaction as predictors, and personal flow score as the outcome measure. We examined this relationship separately for leading, following, and joint improvisation. Overall, spontaneous tapping tempo and SPS did not independently predict flow experience across all conditions. However, their interaction significantly predicted flow (**As Leader**: *β* = −0.65, 95% CI [−1.05, −0.26], *t*(87) = −3.29, *p* = .004, **As Follower**: *β* = −0.56, 95% CI [−0.96, −0.17], *t*(88) = −2.83, *p* = .006, **Joint Improvisation**, *β* = −0.63, 95% CI [−1.05, −0.21], *t*(84) = −2.99, *p* = .005).

To further interpret these interactions, we computed a tempo compatibility index for each participant. First, we standardized spontaneous tapping tempo scores and SPS scores within each condition. We then averaged the standardized SPS scores across the three coordination conditions to extract a single SPS score per participant. To align the directionality of the two measures, we reversed the standardized spontaneous tempo scores, so that relatively fast participants received negative values and slower participants received positive values.

Participants were then grouped based on the alignment between their spontaneous tempo and SPS scores. Those with both scores on the same side of the distribution (i.e., both positive, or both negative) were classified as having high compatibility, indicating that their spontaneous tempo aligned with the temporal structure of joint performance. In contrast, participants with opposing signs (one positive, one negative) were classified as having low compatibility, reflecting a mismatch between their spontaneous tempo and joint performance tempo. We then compared individual flow scores between the two groups using an independent t-test. Individuals in the high compatibility group reported significantly greater flow experience than those in the low compatibility group (mean difference = 5.8, 95% CI [2.10, 9.50], *t* (97) = 3.11, *p* = .002, Cohen’s *d* = 0.63, 95% CI [0.22, 1.04]; see Figure 5). These results suggest that flow experience is enhanced when the temporal structure of joint performance aligns with individuals’ natural tempo tendencies.

## Discussion

In this study, we asked how individuals’ spontaneous tempo shapes joint performance and subjective experience during unstructured, complex and continuous interaction. We found that dyads composed of partners with similar spontaneous tempi reported higher flow experience during joint performance, suggesting that tempo compatibility enhances the quality of shared experience. Surprisingly, we also found that similarity in spontaneous tempo was associated with **reduced** motion synchrony during joint performance. Instead, tempo similarity resulted in a dynamic and unpredictable coordination pattern as reflected by enhanced coordination variability. This increased variability was evident specifically during joint improvisation, where partners could alternate more fluidly between leading and following during the game. Thus, similarity in spontaneous tempi can promote engaging and flexible interaction patterns during joint improvisation.

Furthermore, we asked how individuals’ spontaneous tempo influences the emergent tempo of joint movement. We found that the tempo of joint performance did not reflect individuals’ spontaneous tempo preferences. That is, fast tappers could exhibit a relatively low frequency of motion segments, while slow tappers could produce a relatively high frequency during joint performance. However, the alignment between a participant’s spontaneous tempo and their movement tempo during the interaction significantly impacted their flow experience. Specifically, flow ratings were highest when the tempo of a participant’s motion during the dyadic task corresponded to their spontaneous tempo preferences. This relationship held consistently across all coordination conditions, including leading, following, and joint improvisation. These findings suggest that while individuals can flexibly adapt their movement tempo during social interaction, optimal engagement emerges when the overall tempo of joined movement aligns with their natural motor preferences.

### Spontaneous Tempo Similarity in Interpersonal Coordination

Although several previous studies found that similarity in individuals’ spontaneous movement tempo enhances interpersonal synchrony (Alderisio et al., 2017; Roman et al., 2023; Słowiński et al., 2016; Snapiri et al., 2025, unpublished manuscript; Zamm et al., 2015, 2016), we find that individuals with similar spontaneous tapping tempi synchronize less. This discrepancy can be explained by the fundamentally different characteristics of our task. Previous work focused mostly on highly structured tasks, such as musical performance (Roman et al., 2023; Zamm et al., 2015, 2016), joined rhythmic tasks (Alderisio et al., 2017; Snapiri et al., 2025, unpublished manuscript), or joined movement along a predesignated unidimensional trajectory (Słowiński et al., 2016). Here, we examine joint performance in an unstructured and open-ended task, in which stable synchrony is not necessarily optimal for task performance. On the contrary, several recent works suggest that in complex and dynamic interactions effective coordination often emerges through a continuous interplay between synchrony and breaks from synchrony, allowing for greater creativity and adaptability (Dahan et al., 2016; Laroche et al., 2024; Mayo & Gordon, 2020; Tognoli & Kelso, 2014). Accordingly, we find that while individuals with similar spontaneous tempi overall synchronized less, their coordination pattern was more dynamic and variable. This suggests that in complex and dynamic interactions spontaneous tempo similarity does not lock partners into tight synchrony but instead provides a shared temporal foundation that supports adaptive and expressive coordination (Henry & Kotz, 2023)

### Flow Experience and Coordination Dynamics

Along these lines, we also find that individuals with similar spontaneous tempi reported greater levels of flow during joint performance. This is consistent with recent findings that emphasize the role of novelty and complexity in optimal dyadic experience. For example, Ravreby and colleagues (2022) found that the mutual liking between participants was best predicted by a balance between synchrony and complexity of joint movement, rather than synchrony alone. Similarly, Feniger-schaal and colleagues (2016) found that individuals with secure attachment style favored dynamic exploration over stable synchrony during joint improvisation. And, Wallot and colleagues (2016) found that reduced spontaneous motor synchrony was linked to enhanced subjective experience during a cooperative task in which individuals built a Lego model together, without a pre-designated leader. Taken together, these findings suggest that in complex and creative interactions, coordination variability may better reflect optimal interpersonal engagement rather than overall synchrony.

### Social Distance and Interpersonal Synchrony

The pattern we observed, where individuals with dissimilar spontaneous tempi synchronized **more**, can also be interpreted considering the social functions of motor synchrony. A large body of work suggests that interpersonal synchrony plays an important role in social cohesion and affiliation (Fujiwara et al., 2020; Hove & Risen, 2009; Keller et al., 2014; Marsh et al., 2009; Miles et al., 2011). Miles and colleagues (2011) found that participants synchronized more strongly with members of an out-group than with members of their own social group. They argued that synchrony may emerge as a strategy to reduce perceived social distance and enhance connection. Relatedly, Schmidt and colleagues (1994) found that dyads composed of one partner with high social capacity and one with low social capacity synchronized more than dyads in which both individuals had similar levels of social capacity. These findings have been interpreted in terms of natural reciprocity: in the absence of assigned roles, asymmetries between partners may facilitate the emergence of complementary leader– follower dynamics that support enhanced synchrony. Considering these findings, it is possible that dissimilarity in spontaneous tempo creates a sense of social distance between partners, which in turn motivates them to reduce that distance through increased synchronization. Conversely, when partners are more similar, the drive to reduce distance through synchronization may be weaker, allowing for more exploratory and variable patterns of coordination.

### Tempo Preferences in Joint Performance

An additional aim of the current study was to examine whether and how individuals’ spontaneous tempo influences the temporal structure of joint performance. Prior work has shown that individuals adjust their movement dynamics to facilitate coordination with others. For example, Hart and colleagues (2014) found that during highly synchronized joint action, individuals tend to converge on simplified and predictable motion patterns, minimizing their individual kinematic signatures. Similarly, in a recent study of spontaneous joint tapping, we found that dyads consistently converged to a tapping tempo close to 2 Hz, near the population median for spontaneous tapping tempo (Moelants, 2002). This convergence occurred regardless of the individual spontaneous tempi within each dyad (Snapiri et al., 2025, unpublished manuscript). Therefore, individuals adjusted their spontaneous tempo preferences to support optimal dyadic performance. In a similar manner, in the present study we found that individuals’ spontaneous tempo did not directly predict the movement tempo observed during joint performance. However, when considering the degree of alignment between a participants’ spontaneous tempo and the tempo of their movement during the interaction, a meaningful pattern emerged. Participants reported higher levels of flow when their performance tempo was more closely aligned with their own spontaneous tempo. Therefore, while individuals can flexibly adapt to different tempi of performance, their subjective experience will be affected by the tempo distance they need to traverse to perform together. Future research could further explore how individual tempo preferences shape continuous joint action. For example, whether spontaneous tempo biases emerge asymmetrically across partners or fluctuate over time, contributing to the evolving coordination landscape.

In summary, our findings show that spontaneous tempo can shape both coordination dynamics and subjective experience during unstructured, continuous joint interaction. Tempo similarity between partners did not enhance synchrony but instead supported more dynamic and variable coordination pattern and enhanced flow experience. Additionally, individual flow experience was highest when participants’ dyadic movement tempo aligned with their own spontaneous tempo. These results extend the relevance of spontaneous tempo beyond structured rhythmic tasks, underscoring its role in shaping both behavior and experience in open-ended, dynamic interpersonal coordination.

## Notes

### Competing Interest Statement

The authors have declared no competing interest.

### Funding

- ERC: 101189519, 852387
- Mandel Scholion Research Center

